# Fine-Tuning and Remodelling of Pectins Play a Key Role in the Maintenance of Cell Adhesion

**DOI:** 10.1101/2023.10.27.564335

**Authors:** Cyril Grandjean, Aline Voxeur, Salem Chabout, François Jobert, Laurent Gutierrez, Jérôme Pelloux, Gregory Mouille, Sophie Bouton

## Abstract

Plant cell adhesion is essential for development and stress response, mediated by pectin-rich middle lamella deposition between cell walls. However, the precise control mechanism of cell adhesion remains unclear. The *qua2-1* and *esmd1-1* mutants provide a better understanding of this process and suggest a signaling pathway triggering the loss and restoration of adhesion via cell wall modifications. This study attempts to characterize the potential regulatory role of endogenous oligogalacturonides (OGs) and pectin modifications in the control of cell adhesion in Arabidopsis.

From dark-grown hypocotyls, our extraction revealed seven distinct endogenous OGs with varying polymerization and modifications. Abundance variations of OGs were observed among wild type, *qua2-1*, *esmd1-1*, and *qua2-1/esmd1-1* mutants. The structure of homogalacturonans was analyzed by enzymatic fingerprint, in order to identify changes in esterification patterns. Expression analysis of pectin-modifying enzymes showed significant variations in *PME*, *PMEI*, and *PAE* genes. Gene expressions correlate with homogalacturonans modifications and cell adhesion phenotypes.

This study enhances our understanding of a feedback loop between the endogenous OGs, homogalacturonans esterification fine tuning, and pectin remodeling enzymes expression in controlling cell adhesion.

## INTRODUCTION

Cell adhesion plays a crucial role in plant development and in response to stress (Somerville et al., 2004). The adhesion between adjacent plant cells is facilitated by the deposition of a pectin-rich middle lamella (Daher and Braybrook, 2015). The middle lamella primarily consists of homogalacturonans (HGs), and the degree of methyl and O acetyl esterification of HGs is regulated by cell wall localized proteins such as pectin methylesterases (PMEs), pectin methylesterase inhibitors (PMEIs), and pectin acetylesterases (PAEs) encoded by large multigenic families (Pelloux et al., 2007; Philippe et al., 2017). While some molecular regulators involved in cell adhesion have been identified (Atakhani et al., 2022), the mechanism by which plants control and maintain cell adhesion in response to changes in cell wall chemistry remains poorly understood. Studies on the Arabidopsis *quasimodo2* (*qua2*) mutant, which is affected in a pectin methyltransferase gene and exhibits cellular adhesion defects and reduced amount of HG (Du et al., 2020; Mouille et al., 2007), provided insights into this process. Interestingly, the *esmeralda1* (*esmd1-1*) mutant, carrying a point mutation in a putative *O*-fucosyltransferase, rescues *quasimodo2-1* (*qua2-1*) cell adhesion phenotypes without restoring HG levels (Stéphane Verger et al., 2016). Based on these findings, it has been proposed that *qua2-1* triggers a signal leading to the loss of cell adhesion, and *esmd1* could affect this pathway and restore cell adhesion through cell wall modifications (Stéphane Verger et al., 2016). Identifying the potential signal modified by the *ESMERALDA1* mutation in *qua2-1* and uncovering new actors in this signal transduction pathway will enhance our understanding of the control of cell adhesion in plants.

## RESULTS AND DISCUSSION

### Are endogenous oligogalacturonides (OGs) involved in cell adhesion?

Oligogalacturonides (OGs) are oligomers released from plant cell walls upon partial degradation of homogalacturonans and can activate various signaling pathways when perceived by cell wall receptors (Wolf, 2022). Consequently, OGs are considered as endogenous elicitors that activate plant immunity and control developmental processes (Ferrari et al., 2013; Lin et al., 2022). OGs are the most likely candidates for mediating different signals that could control cell adhesion in Arabidopsis (Stéphane Verger et al., 2016). Therefore, understanding the diversity and roles of endogenous OGs in cell adhesion is essential. In this study, we extracted endogenous OGs from dark-grown hypocotyls and analyzed them by high-performance size exclusion chromatography (HP-SEC) coupled with mass spectrometry (Voxeur et al., 2019). We identified seven different OGs with degrees of polymerization (DP) ranging from 2 to 5, decorated with various methylation and oxidation statuses in the wild type Col-0 and in *qua2-1*, *esmd1-1*, and *qua2-1/esmd1-1* mutants (Figure 1). Although the OG quantities differed among the genotypes, their identities were consistent. Some of these OGs have previously been characterized as potential elicitors, such as the trimer of galacturonic acid (GalA), which triggers a dark-grown signal (Sinclair et al., 2017), and GalA_2_Ox, an OG that accumulates in the later stages of infection by *Botrytis cinerea* (Voxeur et al., 2019). Notably, all the extracted OGs were less abundant in *qua2-1*, with a 50% to 30% decrease compared to the wild type, depending on the OG considered. However, the average quantity of most OGs was partially or fully restored by the *esmd1-1* mutation, and in some cases, the double mutant exhibited even higher levels than the wild type. Statistically, only GalA_4_Me and GalA_4_Me_2_ were significantly more abundant in *esmd1-1* and the double mutant compared to *qua2-1* (Figure 1). The dispersion of values was very low for both mutants. Conversely, there was a wide dispersion of values in the double mutant, as in the wild type, suggesting a recovery of the ability to respond to a stress. As there is no difference in the identity of endogenous OGs, our results suggest there is maybe an involvement of overall amount of these OGs in the recovery of *qua2-1/esmd1-1* cell adhesion.

**Figure 1:**
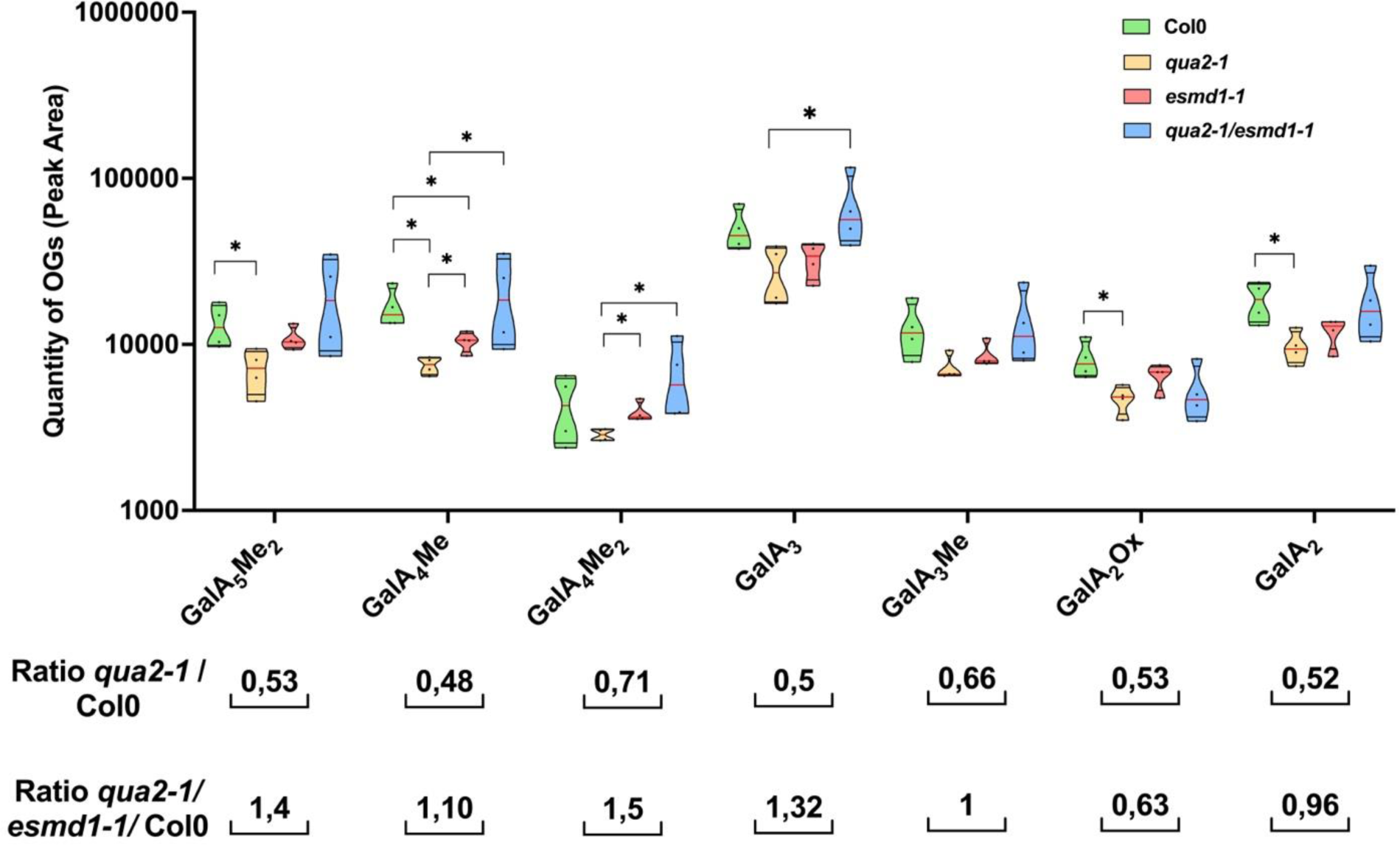
Endogenous OGs. Truncated violin plot of peak area of endogenous OGs of 150 4-day-old dark grown dissected hypocotyls of Col-0, *qua2-1, qua2-1/esmd1-1 and esmd1-1*. Oligogalacturonides (OGs) are named as follow: GalA_x_Me_y_ where the numbers x and y in subscripts indicates the degree of polymerization and the number of methylations respectively. GalA: galacturonic acid; Me: methyl ester group Ox: oxidation. *Red line represents the median, black line the quartiles, black dot represents biological replicate (n = 4 biological replicates per genotype).* *, P < 0.05, Mann & Whitney test.

As pectin digestibility is directly influenced by the esterified pattern and/or the action of various polygalacturonases, pectin and pectate lyases, we further investigated the fine structure of homogalacturonans in the different genetic backgrounds.

### Does the *esmeralda1* mutation alters Homogalacturonans pattern?

To explore the esterification pattern of HGs in different genotypes, we enzymatically digested the cell walls of dark-grown seedlings with a commercial endo-polygalacturonase from *Aspergillus aculeatus*. The released fragments were then analyzed by high-performance size exclusion chromatography (HP-SEC) coupled with mass spectrometry. We identified 92 HG fragments, including monomers with distinct degrees of methyl and O-acetyl esterification, some of which were oxidized. Our results revealed that, independently of the total amount of GalA hydrolyzed by TFA, less HG fragments were released from the total dried cell wall fractions in *qua2-1* compared to the wild type and *esmd1-1* (Supp. Data 1B). Conversely, this digestible fraction of HG, relative to the amount of GalA hydrolyzed by TFA (Figure 2A), remains unchanged in the mutants compared to the wild type, despite a slight decrease being observed. Since the *qua2-1/esmd1-1* double mutant shows a similar pectin defect to *qua2-1* (Verger et al., 2016), this suggests that the fraction of HGs sensitive to PGs digestion, relative to the total amount of HGs, is not the main factor responsible of the restoration of adhesion, as it is more or less maintained across the different genotypes.

**Figure 2.**
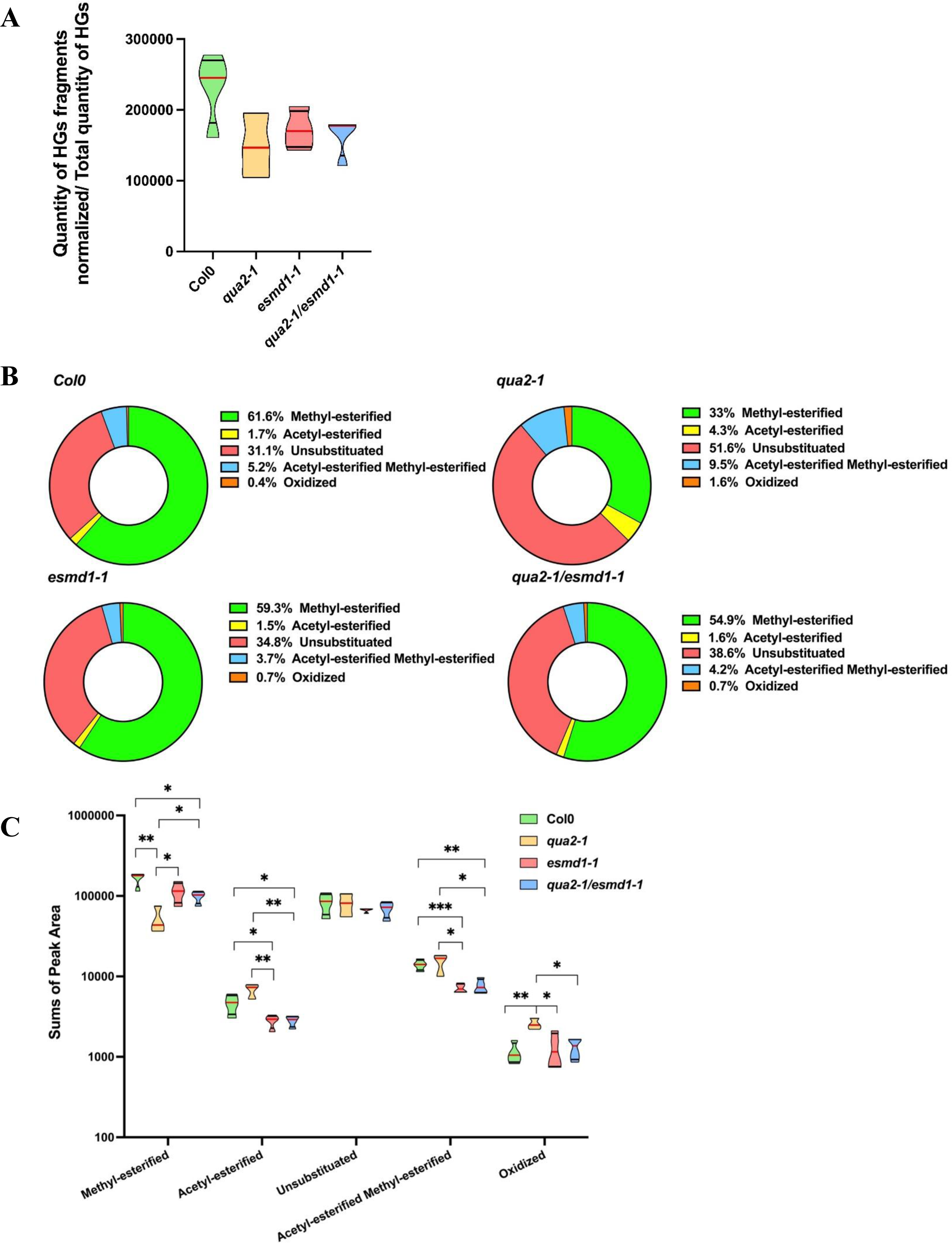
Fine structure of homogalacturonans. (A) Truncated violin plot of the quantity of HGs fragments & monomers released by PG *Aspergillus aculeatus* digestion of dried cell wall seedlings 4 days dark grown of Col-0, *qua2-1, qua2-1/esmd1-1 and esmd1-1*. The quantity was calculated for 1mg of dried cell wall and normalized by the GalA released by TFA hydrolysis. The quantity of GalA hydrolyzed for 1mg of dried cell wall of same samples was analyzed by GC/MS (Supp. data 1A) and subtracted to the equivalent quantity of rhamnose to get the quantity of HGs. HGs fragments were quantified by LC/MS-MS according to Voxeur et al., 2019*. Red line represents the median, black line the quartiles (n ≤ 4 biological replicates per genotype).* *, P < 0.05, **, P < 0.001, t-Test. (B) Relative content of HGs fragments family released by *Aspergillus aculeatus* PG digestion of dried cell wall of 4-days-old dark grown of seedlings Col-0, *qua2-1, qua2-1/esmd1-1 and esmd1-1 (n ≤ 4 biological replicates per genotype).* (C) Truncated violin plot of sums of the quantity of the classes of HGs fragments released by *Aspergillus aculeatus* PG digestion of dried cell wall of 4-days-old dark grown seedlings of Col-0, *qua2-1*, *qua2-1/esmd1-1* and *esmd1-1*. HGs fragments were quantified by LC/MS according to Voxeur et al., 2019. *Red line represents the median, black line the quartiles, (n ≤ 4 biological replicates per genotype).* *, P < 0.05, **, P < 0.001, t-Test.

The identified fragments were grouped into five different categories based on their substitution and oxidation state, reflecting the homogalacturonans pattern. In *qua2-1*, there was a relative decrease (50%) in the amount of methylesterified HGs released compared to the wild type, as previously reported (Du et al., 2020) and to *esmd1-1* (Figure 2B). In contrast, all other fragment categories were 1.5- to 4-fold more abundant in *qua2-1*. This fingerprint analysis revealed a previously unidentified pectin modification in *qua2-1*, indicating a decrease in methyl-esterification of HGs sensitive to polygalacturonase digestion. The proportions of HGs pattern in the *qua2-1/esmd1-1* double mutant suggest that *esmd1-1* restores the relative content of fragment categories, resulting in a proportion of pattern almost indistinguishable from the wild type (Figure 2B), without recovering the HGs quantity (Verger et al., 2016).

Further examination of the quantity of each of the five groups of released HG fragments revealed that the oxidized fragments were more abundant in *qua2-1* compared to the wild type (Figure 2C). In contrast, the group of methyl-esterified fragments were less abundant in *qua2-1*. Except for the oxidized and unsubstituted fragments, the quantities of the other three groups were altered in the *qua2-1*/*esmd1-1* double mutant. It is worth mentioning that the pattern of HG substitution was also altered in the single *esmd1-1* mutant, where the digestion of its cell wall released fewer acetyl and/or methyl-esterified fragments compared to the wild type (Figure 2C).

Taking a closer look at the 49 significantly different HG fragments between the wild type and *qua2-1,* 30 were more abundant in the wild type, while 19 were more abundant in *qua2-1* (Supplementary data 2-A). In the wild type, the accumulated fragments exhibited moderate degree of methyl-esterification (2 to 5) with or without mono-acetyl-esterification, whereas in *qua2-1*, they were highly methyl-esterified (5 to 10), with or without acetyl-esterification (1-3). Thus, the observed modifications in the methyl-esterified pattern *in qua2-1* suggest a defect in PMEs expression or activity.

Analyzing the HG fragments released that were significantly different between *qua2-1* and the double mutant revealed 52 HGs fragments differentially accumulated between the two mutants (Supp.data 2-C). Among them, 13 methyl-esterified (*) and 2 acetyl-esterified (°) and 2 methyl-acetyl-esterified (•) fragments were specific to *esmd1-1*, while 23 methyl-esterified fragments exhibited a similar increase in the wild type compared to *qua2-1* (Supp. data 2-A). Furthermore, 10 methyl-acetyl-esterified, 1 acetyl-esterified and 1 oxidized HG fragments that were significantly more abundant in *qua2-1* compared to the wild type (Supp. data 2-A) remained significantly more abundant in *qua2-1* compared to the double mutant (Supplementary data 2-C). Both *esmd1-1* HGs pattern features, contribute to the restoration of the proportion of methyl-esterified group in *qua2-1* background (Figure 2B). These results also indicate that the *esmd1-1* mutation reduces the acetyl-esterified pattern in the *qua2-1* background, suggesting a higher activity of PAEs that restore the proportion of acetyl-esterified and methyl-acetyl-esterified patterns respectively (Figure 2B). The indirect regulation of HG methyl and O-acetyl esterification patterns by the *O*-fucosyltransferase ESMERALDA is further supported by the comparison between *esmd1-1* and the wild type (Supp.data 2-B).

Overall, since the fraction of HGs sensitive to PGs digestion, relative to the total amount of HGs, remains unchanged, *esmd1-1* in the *qua2-1* background restored the proportion of methyl and/or acetylesterified pattern as well as the oxidized categories of HGs, through a specific modulation of HG pattern including some similarities and divergencies with wild-type. This may lead to a more favorable pattern for proper crosslinking in *qua2*-1 and suggests that pectin pattern modulation allows for more cross-linkable microdomains interspaced within the homogalacturonan domain, enhancing adhesion restoration. It also contributes to the recovery of the overall amount of endogenous OGs (Figure 1).

### Which factors are responsible for changing the pectin pattern of *qua2, qua2/esmd1* and *esmd1*?

To gain a better understanding of how these pectin modifications occur in plants, we performed RT-qPCR analysis of the *PME*, *PMEI* and *PAE* multigenic families in four days-old dark-grown seedlings. Among these genes (respectively 66, 76 and 12), only six exhibited significant expression variations across the genotypes (Figure 3). In *qua2-1*, the expression of *PME53* and *PME41* were increased compared to the wild type. These overexpressions may explain the substantial decrease in methyl-esterified HGs and OGs in the *qua2-1* mutant, which probably leads to the generation of a pectin pattern that is more susceptible to polygalacturonase degradation. These changes in PME gene expression may contribute to the loss of cell adhesion in *quasimodo2*. On the other hand, the expression of *PMEI4* and *PME35* was repressed in the double mutant and *esmd1-1* compared to *qua2-1* (Figure 3). These underexpressions may play a role in the control of cell adhesion by the *esmeralda1* mutation, restoring the relative content of HG methyl-esterified pattern (Figure 2C and Supplementary data 1). Regarding *PAEs*, two genes exhibited similar expression variations. The expressions of *PAE7* and *PAE12* were repressed in *qua2-1* whereas in the double mutant, expressions are similar to the wild type (Figure 3). The restoration of *PAE* expression supports the rebalanced acetyl-esterified and/or methyl-acetyl-esterified patterns in the double mutant (Figure 2C and Supplementary data 2). Acetyl-esterification can prevent HG degradation by endo-polygalacturonases (Bonnin et al., 2003), which may explain the reduction in endogenous OGs in *quasimodo2* (Figure 1).

**Figure 3:**
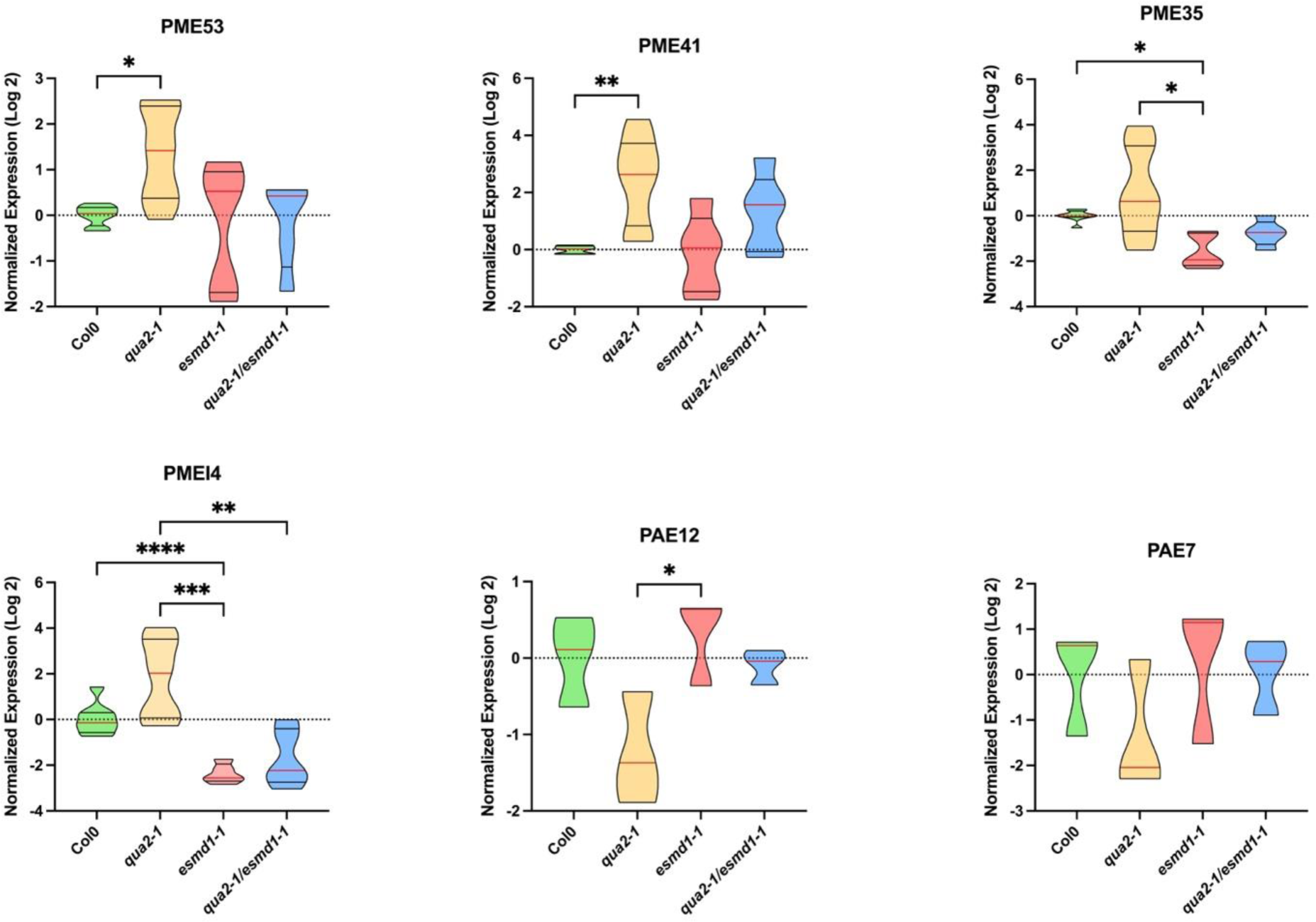
*PME/PMEI* and *PAE* expression. Truncated violin plot of Normalized expression levels (log2) of *PMEI4, PME, 35, 41, 53* and *PAE 7 and 12,* from 4-day-old, etiolated seedlings of Col, *qua2*, *esmd1 and qua2/esmd1* by RT-qPCR. *APT1 & CLATHRIN* were amplified as reference genes. Normalized expression levels (log2) were calculated according to a normalization factor obtained from the 2 reference genes and calibrated to the expression of the wild type (Taylor et al., 2019). *Red line represents the median, black line the quartiles (n ≥ 5 biological replicates per genotype for PME and PMEI) (n = 3 biological replicates per genotype for PAE).* *, P < 0.05 and **, P < 0.001, ***, P < 0.0001, ****, P < 0.00001 Brown-Forsythe and Welch ANOVA tests (Dunn’s multiple comparisons test).

Moreover, the increase in O-acetyl and methyl-O-acetyl-esterified pattern in *qua2-1,* could also compensate in some way the disturbance in the gestion of methyl esterified pattern described above. The transcriptional regulation of *PAE7* and *PAE12* by *esmd1-1* mutation also possibly contributes to the recovery of cell adhesion in *qua2-1*.

Overall, our findings provide evidence of the involvement of PMEs, PMEIs, and PAEs in controlling cell adhesion. As these enzymes belong to large multigenic families, classical reverse genetic approaches often result in compensatory expression, leading to weak phenotypes. However, by focusing on a specific biological phenomenon such as cell adhesion in our case, we were able to better characterize the role of pectin-modifying enzymes. Therefore, we propose that the restoration of cell adhesion in *quasimodo2* by the e*smeralda1-1* mutation is a result of a combination of different mechanisms.

In summary, our findings suggest that the restoration of cell adhesion in *quasimodo2-1* by the *esmeralda1-1* mutation involves the production of endogenous OGs, the fine tuning of HGs pattern and transcriptional modulations of specific pectin-modifying enzymes suggesting a feedback loop regulation (Figure 4). Further investigation into the role of these enzymes and the signaling pathways involved in the process will contribute to a better understanding of the control of cell adhesion in plants.

**Figure 4:**
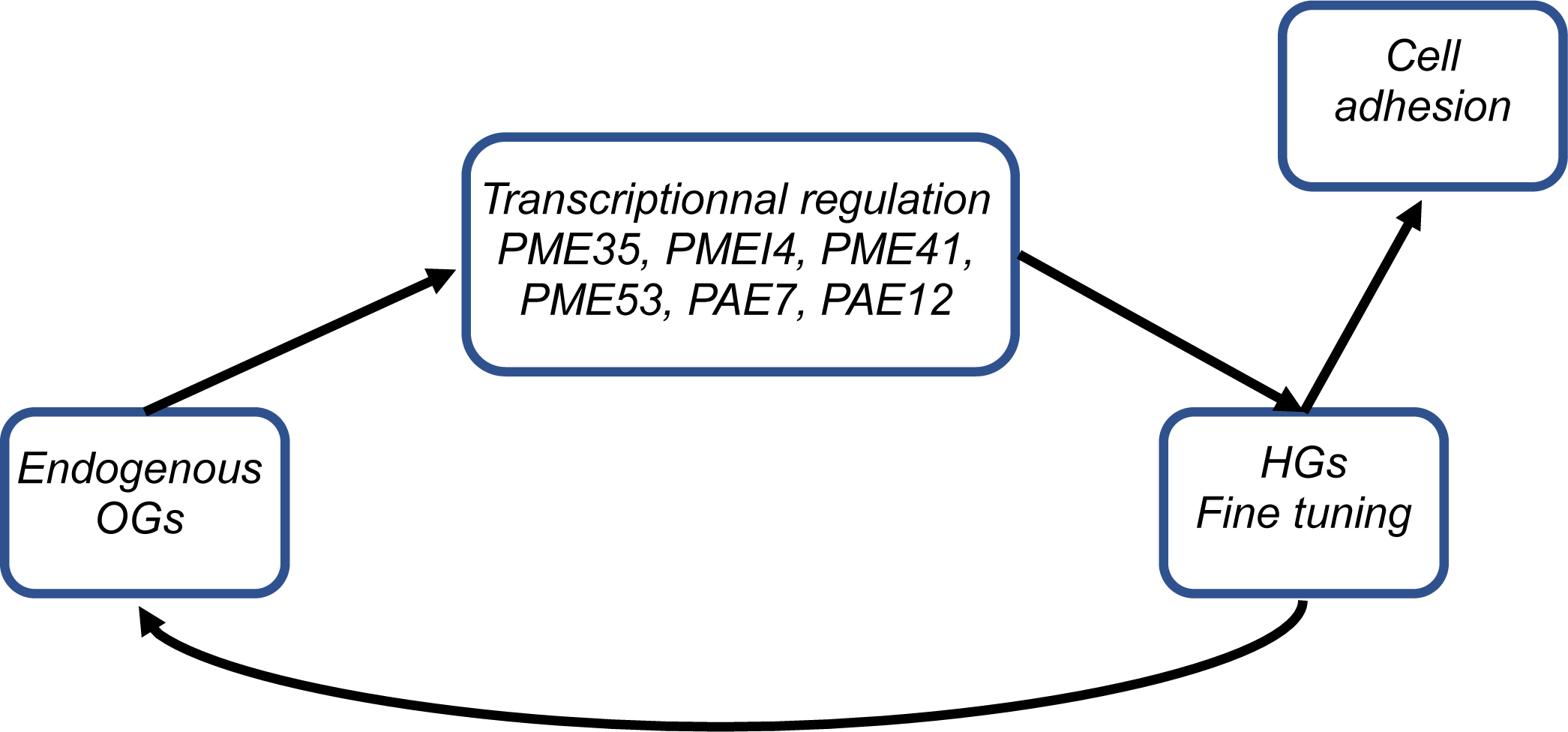
Cell adhesion-pectins feedback loop.

## MATERIALS AND METHODS

### Plant material and growth conditions

Arabidopsis thaliana seedlings (150 seeds/genotype) were grown in the dark at 21 °C on solid medium (Agar 0.8 %, Duchefa Biochemie) supplemented with 0.328 g/L Ca(NO_3_)_2_ at pH 5.7. To synchronize germination, the seeds were cold-treated for 48 hours. After exposure to light for 4 hours, the plates were wrapped in two layers of aluminum foil and cultivated for 92 hours.

### Endogenous OGs extraction

Four biological replicates of 150 four-days-old dark-grown seedlings per genotype were dissected in a dark room to isolate the hypocotyls. The 150 hypocotyls were splitted into 3 tubes (50 in each) and directly frozen. Subsequently, 300 µl of 70 % ethanol were added to each tube. The 3 tubes were then incubated in a thermomixer at 80 °C for one hour. The ethanol-extracted OGs were pooled in the same tube and dried in a speed vacuum concentrator at room temperature. The resulting pellet was re-suspended in 100 µL of ultra-pure water, centrifuged, and the soluble fraction was transferred in a vial and 10 μl was injected for MS analysis.

### Enzymatic fingerprinting of pectins

Four biological replicates of approximately 300 four-days-old dark-grown seedlings were harvested in a dark room and placed in tubes containing 1 ml of 96 % ethanol. The tubes were incubated in a thermomixer at 80 °C for one hour. The previous step with ethanol was repeated for 20 min. The ethanol was then replaced with 1 ml of acetone, and the tubes were incubated in a thermomixer at 25 °C for 20 minutes. This acetone wash step was repeated two more times. The hypocotyls were dried in a speed vacuum concentrator at room temperature overnight, and their dry weight was measured (approximately 1-2 mg per sample). The samples were digested with *Aspergillus aculeatus* endo-polygalacturonase M2 (Megazyme) in 50 mM ammonium acetate buffer pH 5 at 37 °C for 18 hours. After inactivating the enzyme by heating, the digested fractions were collected for MS analysis.

### OGs characterization and quantification by LC/HRMS analysis

The endogenous OGs and the OGs released from the digestion were analyzed using High-performance size-exclusion chromatography (HP-SEC) coupled with MS. The chromatographic separation was performed on an ACQUITY UPLC Protein BEH SEC Column (125 Å, 1.7 μm, 4.6 mm x 300 mm) at a flow rate of 400 µl/min and a column oven temperature of 40 °C. The MS detection was performed in negative mode with the parameters described in the supplementary Materials and Methods.

### RNA extraction and RT-qPCR analysis

Five biological replicates of approximately 150 seedlings grown in the dark for four days were harvested in a dark room. Total RNA was isolated using the RNeasy Plant Mini Kit (Qiagen) with on-column DNA digestion using Rnase-Free DNase (Qiagen). Two µg of total RNAs were used to synthesize cDNAs using RevertAid H minus reverse transcriptase, according to the manufacturer’s instructions Thermofisher.

Transcript levels were assessed by quantitative RT-PCR using a Light Cycler® 480 System (ROCHE), as previously described by (Gutierrez et al., 2009). The expression levels of the target genes were normalized to the reference genes *CLATHRIN* (At5g46630) and *APT1* (At1g27450), which were selected with GENORM (Vandesompele et al., 2002), according to the method described in (Taylor et al., 2019). The primer sequences for RT-qPCR are provided in the supplementary Materials and Methods.

**Table 1.**
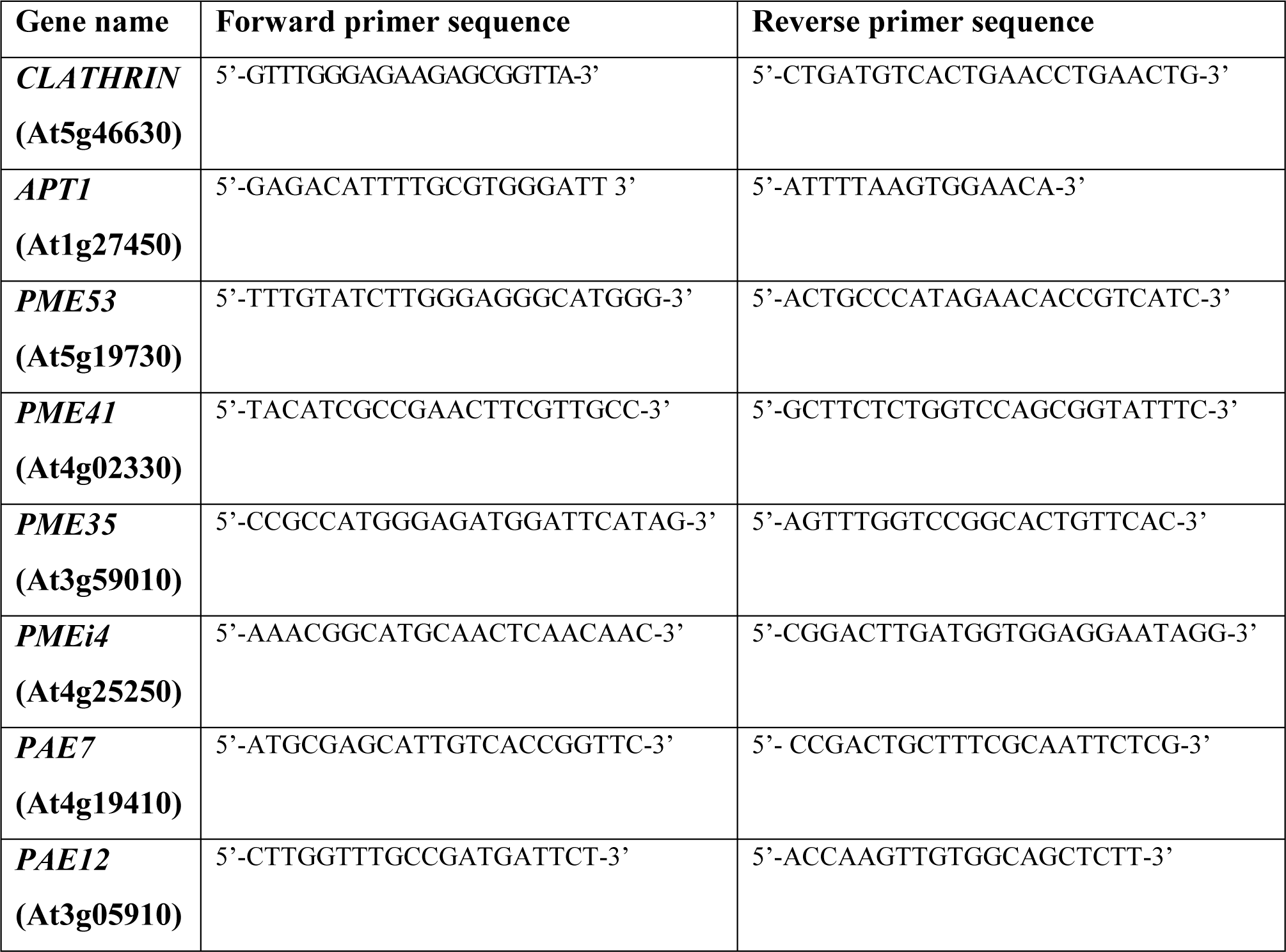
List of primers used for qPCR.

## Supplementary Material and Methods

### Monosaccharides quantification by GC/MS

The monosaccharides of the cell wall were analyzed with the protocol according to (Clement et al., 2018). Four biological replicates of approximately 300 seedlings cultivated in the dark over 4 days were harvested in a dark room and placed directly in the 96 ° ethanol for 1 h at 80 ° C to fix the sample. After removing the ethanol, the first step is repeated for 20 minutes. 1 ml of acetone is added and the tubes are placed in a thermomixer for 20 min at 25 ° C, this step is repeated twice. The cell walls are then dried in a high vacuum concentrator at room temperature overnight. 1 mg dry cell wall previously weighed is resuspended in 400 L of TFA (2 M, freshly prepared), and incubate at 120 ° C for 1 h in a heat block in 1.5 mL screwcap tubes. Then samples were centrifugated for 10 min. The supernatant was transferred and dried in a speed vacuum concentrator at room temperature.

#### Derivatization

after adding 10 μl of 20 mg/ml methoxamine in pyridine to the samples, the reaction was performed for 90 min at 28 °C under continuous shaking in an Eppendorf thermomixer. 50 μl of N-methyl-N-trimethylsilyl-trifluoroacetamide (MSTFA) (Sigma M7891-10×1 mL) were then added and the reaction continued for 30 min at 37 °C. After cooling, 45 μl were transferred to an Agilent vial for injection.

#### Data processing

Raw Agilent datafiles were converted in NetCDF format and analyzed with AMDIS http://chemdata.nist.gov/mass-spc/amdis/. A home retention indices/ mass spectra library built from the NIST, Golm, http://gmd.mpimp-golm.mpg.de/ and Fiehn databases and standard compounds were used for metabolites identification. Peak areas were also determined with the Targetlynx software (Waters) after conversion of the NetCDF file in masslynx format. AMDIS, Target Lynx in splitless and split 30 modes were compiled in one single Excel file for comparison. After blank mean subtraction, peak areas were normalized to Ribitol and dry weight.

#### Absolute quantification

A response coefficient was determined for 4 ng each of a set of monosaccharides, respectively to the same amount of ribitol. This factor was used to give an estimation of the absolute concentration of the metabolite in what we may call a “one point calibration”.

High-performance size exclusion chromatography (HP-SEC) coupled with mass spectrometry analysisChromatographic separation was conducted using an ACQUITY UPLC Protein BEH SEC Column (125Å, 1.7 μm, 4.6 mm × 300 mm, Waters Corporation, Milford, MA, USA) coupled with a guard Column BEH SEC Column (125Å, 1.7 μm, 4.6 mm × 30 mm). Elution was performed with a flow rate of 400 µl/min at a column oven temperature of 40 °C using 50 mM ammonium formate containing 0.1 % formic acid. The injection volume was set to 10 µl. ESI MS-detection was carried out in negative mode on a Bruker impact II QTOF with the following parameters: end plate offset set voltage of 500 V, capillary voltage of 4000 V, Nebulizer at 40 psi, dry gas flow at 8 l/min, and dry temperature at 180 °C. Data acquisition was performed using Compass 1.8 software (Bruker Daltonics). Data analysis was conducted using Mzmine 2.53 software, as described by Pluskal et al. (2010). The integration process involved the following steps: mass detection with a filter noise level set to 500, ADAP Chromatogram Builder using the parameters range of 6 - 9.60 min, minimum group size of scan of 10, group intensity threshold of 1500, minimum highest peak of 1000, and m/z tolerance of 0.01 or 10 ppm. The chromatogram peaks were deconvoluted using baseline cut-off with the baseline level set at 300. Chromatograms were deisotoped with an m/z tolerance of 0.01 or 5 ppm, retention time tolerance of 0.1, maximum charge of 2, and representative isotope as the most intense. Subsequently, peaks were aligned with an m/z tolerance of 0.01 or 5 ppm, retention time tolerance of 0.1, and equal weight for m/z and retention time (weight=1). Finally, a gap-filled peak finder was applied with an m/z tolerance of 0.01 or 5 ppm, retention time tolerance of 0.1, and intensity tolerance of 20 %. The peak areas were exported at the end.

**Supplemental data 1 :**
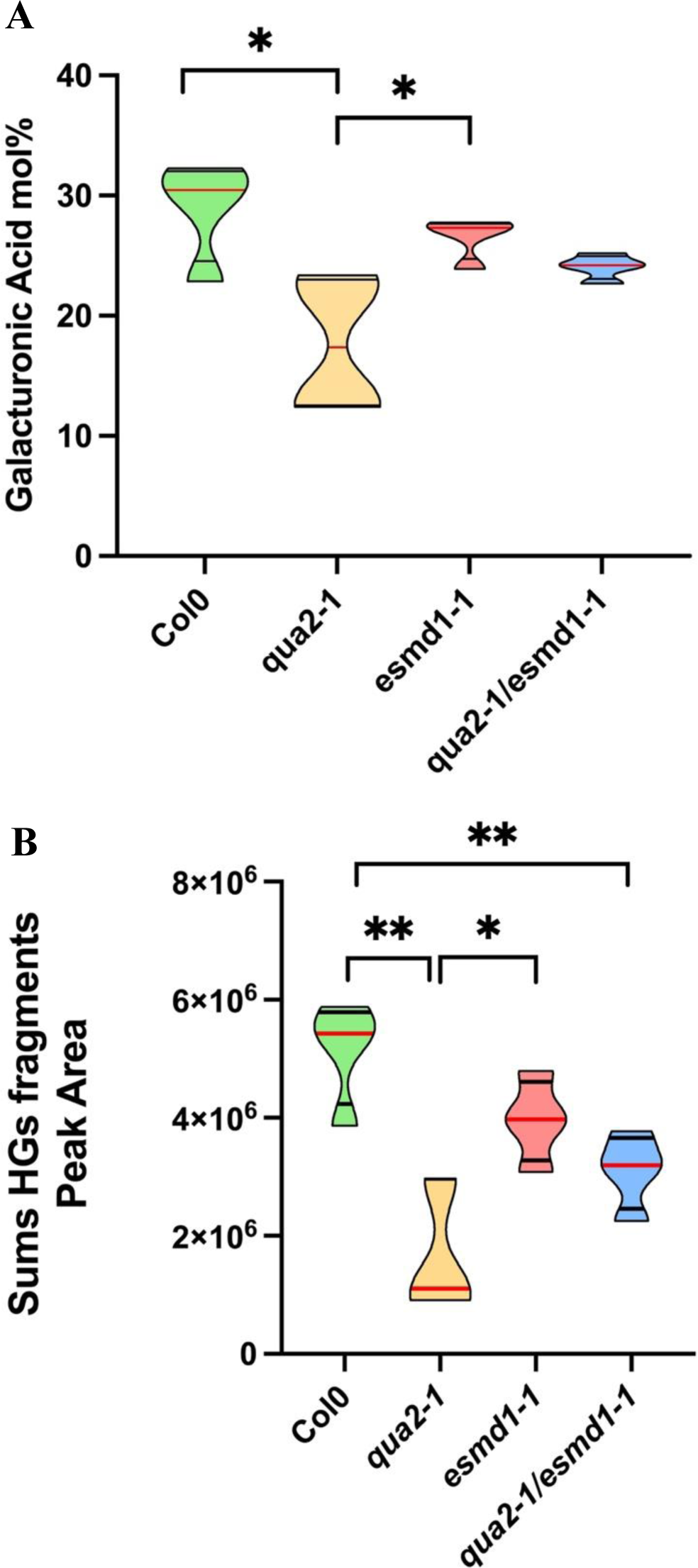
Galacturonic acid content and sums of HGs fragments released. (A)Truncated violin plot of Galacturonic acid content relative to the rhamnose content, represented in mol% and quantified by GC/MS on dried cell wall seedlings 4 days dark grown of Col-0, *qua2-1, qua2-1/esmd1-1 and esmd1-1. Red line represents the median, black line the quartiles (n =4 biological replicates per genotype).* *, P < 0.05, T-Test. (B)Truncated violin plot of the quantity of HGs fragments & monomers released by PG *Aspergillus aculeatus* digestion of dried cell wall seedlings 4 days dark grown of Col-0, *qua2-1, qua2-1/esmd1-1 and esmd1-1*. *, P < 0.05, **, P < 0.001, T-Test

**Supplemental data 2 :**
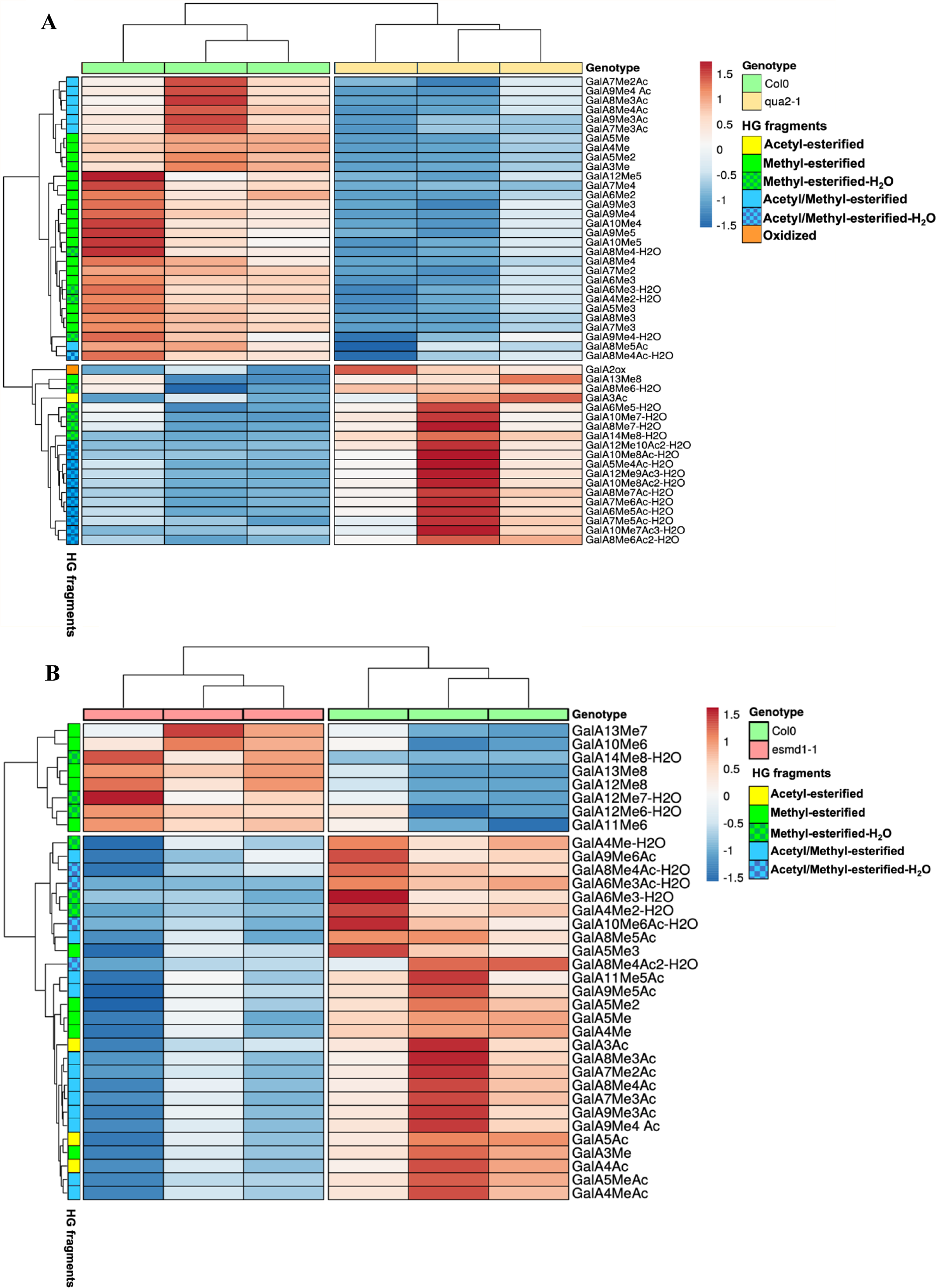
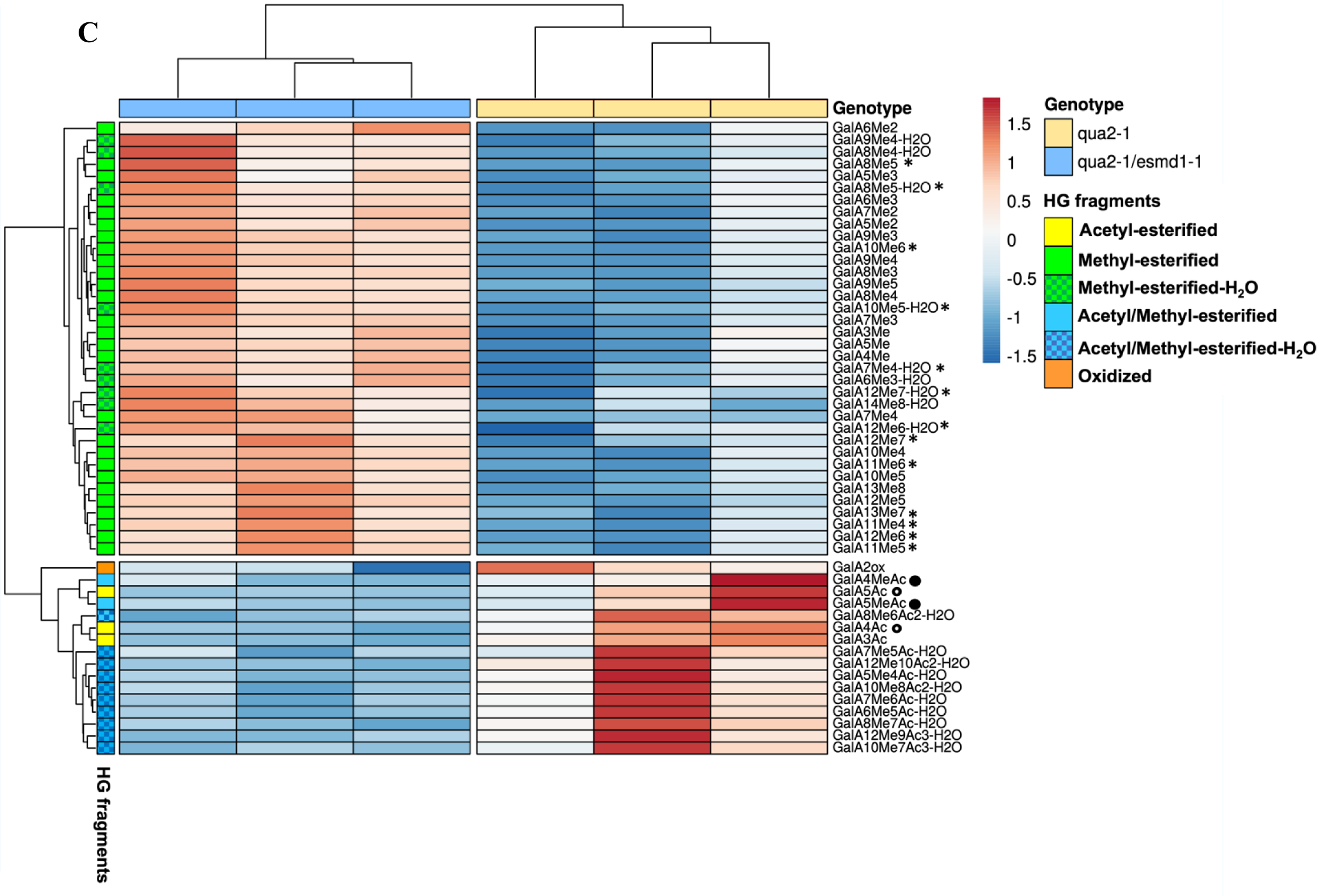
HGs fragments released significantly different. (A) Heatmap of all of the HGs fragments released by *PG Aspergillus aculeatus* significantly different between Col0 and *qua2-1* (46 rows) for 3 samples (columns). Annotation labels refer to HGs fragments structure. Rows are centered; unit variance scaling is applied to rows. Both rows and columns are clustered using McQuitty distance and maximum linkage (B) Heatmap of all of the HGs fragments released by *PG Aspergillus aculeatus* significantly different between Col0 and *esmd1-1* (35 rows) for 3 samples (columns) (C) Heatmap of all of the HGs fragments released by *PG Aspergillus aculeatus* significantly different between *qua2-1* and *qua2-1/esmd1-1* (53 rows) for 3 samples (columns). Methyl- esterified (*) and acetyl-esterified (°) methyl/acetyl-esterified (•) HGs fragments specific *to esmd1-1* are highlighted. For (A), (B), and (C) *(n = 3 biological replicates per genotype),*

## Acknowledgements

The IJPB benefits from the support of Saclay Plant Sciences-SPS (ANR-17-EUR-0007).

## Author contributions

S.B., G.M. and J.P. conceived the research project. S.B., G.M. and C.G. conceived and designed the experiments. C.G. performed most of the experiments with input from S.C. and A.V. L.G. and F.J. contribute to transcriptomic experiments. S.B., GM and C.G. wrote the manuscript.

## References

Atakhani, A., Bogdziewiez, L. and Verger, S. (2022). Characterising the mechanics of cell– cell adhesion in plants. Quant. Plant Biol. 3,.

Bonnin, E., Le Goff, A., Van Alebeek, G. J. W. M., Voragen, A. G. J. and Thibault, J. F. (2003). Mode of action of Fusarium moniliforme endopolygalacturonase towards acetylated pectin. Carbohydr. Polym. 52, 381–388.

Daher, F. B. and Braybrook, S. A. (2015). How to let go: Pectin and plant cell adhesion. Front. Plant Sci. 6, 1–8.

Du, J., Kirui, A., Huang, S., Wang, L., Barnes, W. J., Kiemle, S. N., Zheng, Y., Rui, Y., Ruan, M., Qi, S., et al. (2020). Mutations in the Pectin Methyltransferase QUASIMODO2 Influence Cellulose Biosynthesis and Wall Integrity in Arabidopsis [OPEN]. Plant Cell 32, 3576–3597.

Ferrari, S., Savatin, D. V., Sicilia, F., Gramegna, G., Cervone, F. and De Lorenzo, G. (2013). Oligogalacturonides: Plant damage-associated molecular patterns and regulators of growth and development. Front. Plant Sci. 4, 1–9.

Gutierrez, L., Bussell, J. D., Pǎcurar, D. I., Schwambach, J., Pǎcurar, M. and Bellini, C. (2009). Phenotypic plasticity of adventitious rooting in arabidopsis is controlled by complex regulation of AUXIN RESPONSE FACTOR transcripts and microRNA abundance. Plant Cell 21, 3119–3132.

Lin, W., Tang, W., Pan, X., Huang, A., Gao, X., Anderson, C. T. and Yang, Z. (2022). Arabidopsis pavement cell morphogenesis requires FERONIA binding to pectin for activation of ROP GTPase signaling. Curr. Biol. 32, 497–507.e4.

Mouille, G., Ralet, M. C., Cavelier, C., Eland, C., Effroy, D., Hématy, K., McCartney, L., Truong, H. N., Gaudon, V., Thibault, J. F., et al. (2007). Homogalacturonan synthesis in Arabidopsis thaliana requires a Golgi-localized protein with a putative methyltransferase domain. Plant J. 50, 605–614.

Pelloux, J., Rustérucci, C. and Mellerowicz, E. J. (2007). New insights into pectin methylesterase structure and function. Trends Plant Sci. 12, 267–277.

Philippe, F., Pelloux, J. and Rayon, C. (2017). Plant pectin acetylesterase structure and function: New insights from bioinformatic analysis. BMC Genomics 18, 1–18.

Sinclair, S. A., Larue, C., Bonk, L., Khan, A., Castillo-Michel, H., Stein, R. J., Grolimund, D., Begerow, D., Neumann, U., Haydon, M. J., et al. (2017). Etiolated Seedling Development Requires Repression of Photomorphogenesis by a Small Cell-Wall-Derived Dark Signal. Curr. Biol. 27, 3403–3418.e7.

Somerville, C., Bauer, S., Brininstool, G., Facette, M., Hamann, T., Milne, J., Osborne, E., Paredez, A., Persson, S., Raab, T., et al. (2004). Toward a Systems Approach toUnderstanding Plant Cell Walls. Science (80-.). 306, 2206–2211.

Stéphane Verger, S., Chabout, S., Gineau, E. and Grégory Mouille, G. (2016). Cell adhesion in plants is under the control of putative O-fucosyltransferases.

Taylor, S. C., Nadeau, K., Abbasi, M., Lachance, C., Nguyen, M. and Fenrich, J. (2019). The Ultimate qPCR Experiment: Producing Publication Quality, Reproducible Data the First Time. Trends Biotechnol. 37, 761–774.

Vandesompele, J., De Preter, K., Pattyn, F., Poppe, B., Van Roy, N., De Paepe, A. and Speleman, F. (2002). Accurate normalization of real-time quantitative RT-PCR data by geometric averaging of multiple internal control genes. Genome Biol. 3,.

Voxeur, A., Habrylo, O., Guénin, S., Miart, F., Soulié, M. C., Rihouey, C., Pau-Roblot, C., Domon, J. M., Gutierrez, L., Pelloux, J., et al. (2019). Oligogalacturonide production upon Arabidopsis thaliana-Botrytis cinerea interaction. Proc. Natl. Acad. Sci. U. S. A. 116, 19743–19752.

Wolf, S. (2022). Cell Wall Signaling in Plant Development and Defense. Annu. Rev. Plant Biol. 73, 323–353.

